# Increasing the effective gene drive homing rate by targeting the haploinsufficient spermatogenesis gene *KLHL10*

**DOI:** 10.64898/2025.12.22.695236

**Authors:** Ceili L. Peng, W. Sebastian Kamau, Julien Freeman, Zachary J. Hill, Kevin M. Esvelt

## Abstract

CRISPR-based gene drives represent a powerful new technology for limiting disease transmission and controlling invasive populations. These systems rely on homology-directed repair (HDR) to ‘drive’ a genetic element through a population. However, mammals tend to favor non-homologous end joining (NHEJ), which generates mutations that halt further drive propagation. Here, we describe the experimental characterization of a gene drive system targeting the haploinsufficient spermatogenesis gene *KLHL10* in the laboratory mouse. Using a newly designed ‘coding sequence cassette’ we introduce downstream guide RNAs within the gene, ensuring that sperm undergoing NHEJ are selectively removed from the population. As a proof of principle, we demonstrate that targeting KLHL10 with constitutively expressed LbCas12a results in strong selection against frameshift-containing sperm, validating the core purification mechanism required for this drive strategy. Unexpectedly, we also observed that female offspring lacked most frameshift mutations, suggesting a previously unrecognized role for *KLHL10* in oogenesis or early embryonic development.

## Introduction

Engineered gene drive is a powerful genetic technology designed to propagate specific genetic traits through populations at rates exceeding Mendelian inheritance. First conceptualized in the early 2000s and experimentally realized through CRISPR-Cas systems in the 2010s, most gene drive systems function by converting germline cells in heterozygous individuals into homozygotes for the drive allele, resulting in preferential transmission to offspring^1–4^. This “super-Mendelian” inheritance has profound implications for ecological management, disease control, and evolutionary biology.

In wild populations, gene drives could control disease vectors like malaria-transmitting mosquitoes or preserve endangered organisms through genetic rescue, while localized forms might directly control populations of invasive species or pests^2,5,6^. Initial successes in Anopheline mosquitos demonstrated high transmission rates exceeding 95% in both sexes in some laboratory studies^7^. However, the efficiency of gene drives varies considerably across species, with technical implementation challenges including localization, the evolution of resistance alleles, and importantly, differences in DNA repair pathway preferences^8^. The latter challenge is particularly pronounced in mammals, where it has proven difficult to bias double-strand break repair towards homology-directed repair (HDR) compared to non-homologous end joining (NHEJ)^9–11^. While HDR faithfully copies the drive element, NHEJ often creates indels that render the drive non-functional and generate resistance alleles.

The highest reported rates of HDR gene conversion in *Mus musculus*, 72% in females and 11% in males, are likely too low for efficient gene drive^9,10^. In *Rattus norvegicus*, the highest reported rates of reusable HDR gene conversion are 67% in females and 0.9% in males^12^. This strong preference for NHEJ in rodents, especially in males, has severely limited gene drive efficiency in these organisms(depicted in Fig. 1A).

**Fig. 1:**
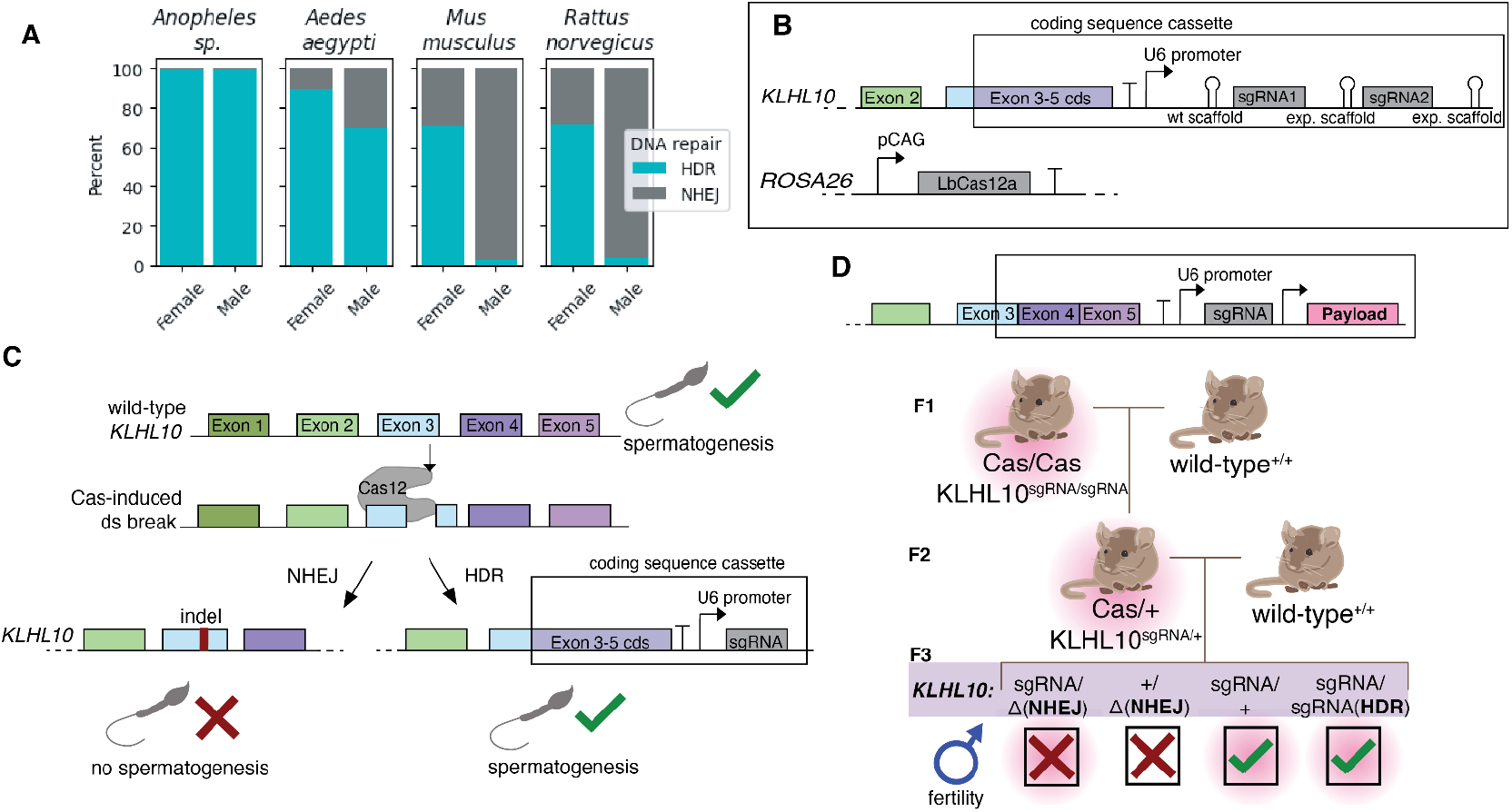
A proposed system of gene drive to compensate for low rates of HDR in males. A.) Previous research demonstrates that rates of homology directed repair are low in mammals, but especially low in males^9,10,12,14,15^. B.) Our split-drive system consists of a ‘coding sequence cassette’ (cds) that supplements the remaining protein coding sequence of the *KLHL10* gene followed by an array of LbCas12a guides/scaffolds. LbCas12a was expressed out of the *ROSA26* safe harbor locus. C.) A gene drive targeting a haploinsufficient spermatogenesis gene: LbCas12a first cleaves an exon of the target gene, then the double-strand break is either repaired by NHEJ, resulting in gene knockout, or by HDR, which rescues the function of the gene by inserting the cds and a downstream guide RNA. D.) Operationalization: By delivering a genetic payload alongside the guide RNA, one could drive the payload through a population of wild-type mice. Mice containing the ‘payload’ are denoted in pink.

Our approach examines a potential solution to the low gene drive homing efficiency in male mice by targeting *KLHL10*, a gene that reportedly exhibits haploinsufficiency for spermatogenesis in mice^13^. Rather than attempting to increase HDR rates directly, we exploit the natural consequences of NHEJ at this locus: frameshift mutations render sperm non-functional, effectively ensuring that only drive-carrying or uncut sperm can produce offspring. This strategy eliminates NHEJ alleles at the abundant sperm stage, effectively decreasing the inheritance rate of mutated, drive-resistant alleles.

Our experimental system targets *KLHL10* using the RR-variant of the Lachnospiraceae bacterium (Lb) variant of Cas12a (LbCas12a), which we selected for its multiplexing potential and targeting versatility. To investigate its utility, we generated transgenic mice expressing LbCas12a and guide RNAs targeting the *KLHL10* locus in a split-drive system. Our analysis focused on understanding LbCas12 cutting efficiency, mutation patterns, and the feasibility of using this system for gene drive applications.

## Results

### A system of drive targeting haploinsufficient spermatogenesis genes

Our design targets the *KLHL10* gene, reportedly haploinsufficient for spermatogenesis^13^, with the LbCas12a(RR) endonuclease to overcome low mammalian HDR rates through strong selection against NHEJ-containing sperm (Fig. 1A). Because initial attempts to express LbCas12a from the meiosis-specific *SPO11* locus did not yield appreciable cutting rates (data not shown), our experimental design tested the feasibility of sperm-specific selection for HDR by constitutively expressing LbCas12a with the CAG promoter from known safe harbor locus *ROSA26* (Fig. 1B). Given that rates of HDR in mammals are optimized using meiotic germline-specific promoters, we did not expect to see instances of HDR in our study, rather, we sought to experimentally test the efficacy of targeting spermatogenesis gene *KLHL10* with CRISPR to select against alleles modified by NHEJ.

As previous research has shown that male mice lacking two functioning alleles of *KLHL10* are infertile^13^; we hypothesized that in the context of a drive system NHEJ repair will produce indels that will disrupt spermatogenesis, while HDR introduces a ‘coding sequence cassette’ containing the remaining protein sequence needed for *KLHL10* function (Fig. 1C). Importantly, this system is amenable to driving a genetic payload downstream of the guide targeting the spermatogenesis gene. The payload (Fig.1D) could be another guide RNA or a protein such as an antibody. In order to test a design with multiple guide RNAs, which can increase genetic stability, we evaluated a dual-guide system incorporating a wild-type crRNA scaffold with a guide RNA targeting exon 3, and an experimental scaffold with a guide RNA targeting a downstream site in exon 3 (Fig. 1B).

### LbCas12a in mice

To robustly measure the *in vivo* cutting efficiency of LbCas12a, we analyzed the genetics of *KLHL10* in F2 mice (Fig. 2) by performing targeted Sanger sequencing followed by trace decomposition analysis of ear tissue samples^16^. Out of 21 initial offspring, 8 (38%) exhibited detectable mosaicism (Fig. 2B). Of the 8 mosaic mice, edits were observed in 4% of cells to 87% of cells, with an average of 36% of cells containing indels (Fig. 2C). We were able to detect a wide range of indels, from -17 to +14 (Fig. 3A).

**Fig. 2:**
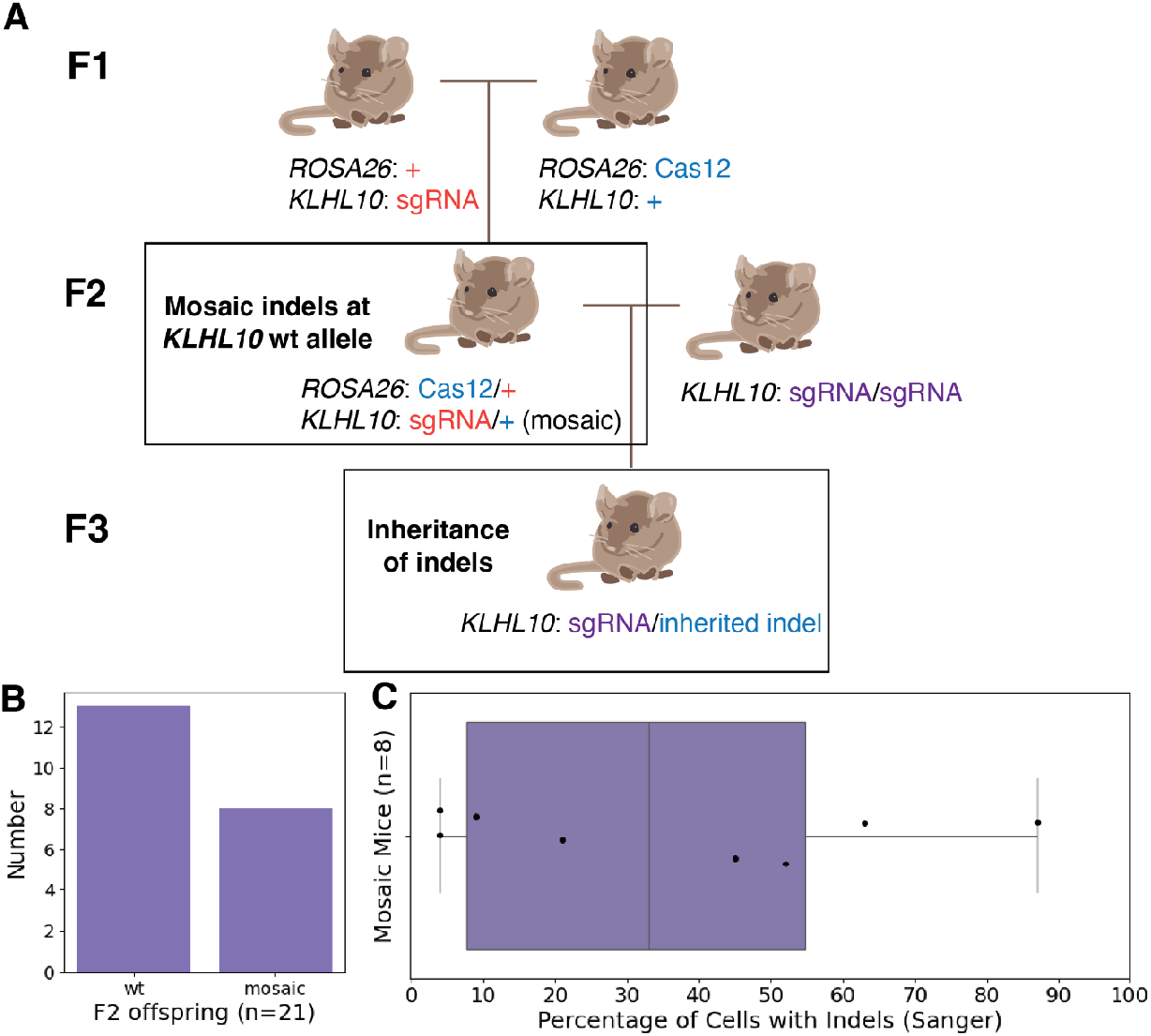
LbCas12a has moderate, varied cutting efficiency *in vivo*. A.) Experimental breeding scheme. F2 mice were analyzed for mosaicism at the ‘wt’ *KLHL10* locus, F3 mice heterozygotic for the engineered allele were also genotyped at the ‘wt’ locus to confirm inheritance of indels. B.) F2 offspring containing mosaicism at the intended cut site vs offspring where LbCas12a did not produce any genetic edits. C.) Percentage of cells within individual F2 mosaic mice containing indels.

**Fig. 3:**
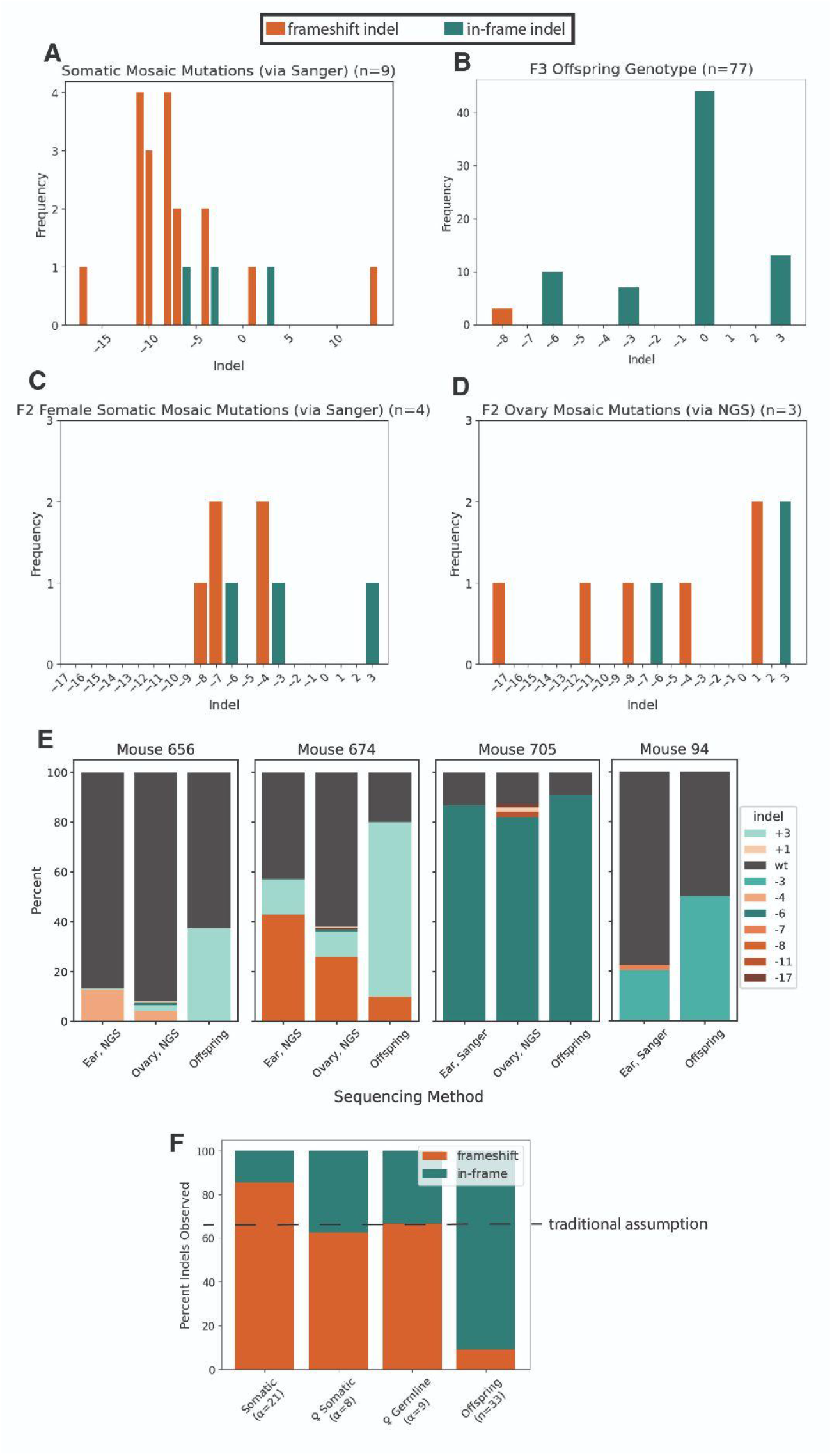
Selection for in-frame mutations purifies *KLHL10* alleles. A) Frequency of somatic indel mutations observed in mosaic mice (prevalence within each mouse’s tissues not included). Includes one mosaic mouse (946) present in the F3 generation that did not inherit the mutation from its parent, but not a mosaic mouse that did inherit mutations (988). B) Frequency inherited indels in F3 offspring. C.) Occurrence of indel mutations in mosaic female mice that produced F3 offspring, measured via Sanger and trace decomposition analysis. D) Indel occurrence in ovaries of three F2 female mice that produced F3 offspring, measured via NGS^17^. E) Prevalence of different somatic and germline indel mutations in female F2 mice that produced F3 offspring, compared to incidence in those offspring. F) Proportion of frameshift and in-frame mutations observed in different genotyped cohorts. Traditional assumption denotes 66% frameshift indels. (n = number of mice, α = number of observed indel events)

Male mice with mosaic edits either produced offspring without indels (n=2) or no offspring (n=1). This is consistent with previous research that found that low percentage *KLHL10* chimeric males were able to produce wildtype offspring, but that high percentage chimeric males were infertile ^13^. Notably, in our experiment, males with a lower frequency of mosaicism (4% and 50% somatic tissue mosaicism) produced offspring, whereas the infertile male featured 63% knockout mutations in somatic tissue and 91% in germline tissue (Supplemental Table 1). In establishing and maintaining the mouse line, we observed that male mice that were heterozygotic or homozygotic for this knock-in cassette were able to produce typical numbers of offspring that successfully inherited our engineered allele, confirming that the ‘coding sequence cassette’ restores function of the gene and that there are no obvious fitness defects associated with our design (Supplemental Table 1). Given the constitutive CRISPR activity outside the meiotic window, we predictably did not observe homology directed repair.

Female mosaic mice produced typical numbers of offspring (Supplemental Table 1). However, we were surprised to see that virtually all of these offspring contained non-frameshift mutations (Fig. 3B). Given that our initial Sanger sequencing of the mosaic parents yielded a wide variety of edits (Fig. 3A), we investigated the consequences of *KLHL10* mutations for viability. We first confirmed that our analysis was accurate using NGS; we found similar results from the trace decomposition analysis and NGS/CRISPResso2 analysis^16,17^ (Supplemental Figure 1). We then dissected mice to compare the somatic and germline tissues, given that various tissues can have different ratios or kinds of indels, to explore the possibility of previously undescribed developmental functions of *KLHL10* (Fig. 3C,D,E).

### *KLHL10* Gene Analysis

Initial analysis indicated that 86% of observed mutations in parent somatic tissue were frameshifts, whereas only 9% of offspring inherited frameshift mutations(Fig. 3F). In some breeding pairs (mouse 656 and 674), a mutation that was a minority mutation in the somatic and germline tissue was heavily favored in offspring, suggesting some form of selection (Fig. 3E).

The only frameshift mutation that was inherited by offspring mice was a single -8 deletion. Given the assumption that ⅔ of mutations will be frameshift mutations and ⅓ of mutations will be in-frame mutations in the absence of selection, we calculated the probability of this occurrence using a cumulative binomial distribution. We observed 33 mice that inherited indels from 4 mosaic mothers; of those, 30 of them inherited in-frame mutations (+3, -3, -6), whereas 3 inherited a frameshift mutation (-8). Of 33 offspring, where P(frameshift) = 2/3, the cumulative binomial probability of 3 or fewer mice containing frameshift mutations by chance is 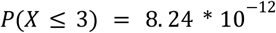(Fig. 3F).

To ensure that our results were not biased by differential somatic and germline editing, we performed a similar analysis on the distribution of indels present in the parent mouse ovarian tissue (Fig. 3D). There were 9 types of indels detected in the ovaries of 3 females, of which 3 were in-frame mutations, suggesting that there is no selection against somatic editing (Fig. 3F). However, these same females produced 23 in-frame offspring and 3 frameshift offspring; therefore, given 26 offspring and P(frameshift) = (2/3), the cumulative binomial probability of 3 or fewer such offspring is 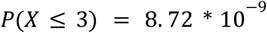(Fig. 3F). Given this consistently improbable result, our data suggest that the *KLHL10* gene may have some previously undescribed function in female fertility or embryo development in addition to its described role in spermatogenesis.

We went on to successfully breed a heterozygote female mouse with the -8 deletion, observing no inheritance bias (Supplemental Figure 2B). We similarly bred F3 heterozygote males and females with in-frame mutations (+3 and -6), and did not observe detectable inheritance biases between the mutations and the engineered allele (Supplemental Figure 2A). Therefore, our targeted mutation site purifies most frameshift mutations in both males and females, but not the -8 frameshift or the observed in-frame mutations. As there is no crystal structure of the *KLHL10* protein, we used AlphaFold to examine the predicted structure and visualize our mutation target^18^ (Supplemental Figure 3A). All indels occur in an exterior loop, and none of them are predicted to alter the overall structure of the protein (Supplemental Figure 3B). The reason that the -8 frameshift mutation does not impair female development, unlike all other possible frameshifts, is unclear.

### Modeling Spermatogenic selection

To evaluate the population-level dynamics of our purifying gene drive system, we adapted a previous modeling framework for mouse populations^19^. We parameterized the model using the sex-specific germline activity observed in females in Grunwald et al (2019) and in males in Weitzel et al (2021): female homing efficiency of 37.6% (calculated as pC(1-pN) where pC=0.552 and pN=0.319) and a male homing efficiency of 3.0% (pC=0.289, pN=0.895)^9,10^. The model simulated a population of 10,000 mice following a one-time release of 100 D/D homozygous drive-carrying individuals. Purification is modeled as occurring during germline activity in each breeding cycle in both female and male drive carriers. When drive-carrying individuals produce gametes, NHEJ-derived resistance alleles (r2) are subject to purification with a specified efficiency. The remaining gametes are then renormalized, effectively increasing the transmission bias toward drive and wild-type alleles.

In the absence of purification (Fig.4A), both the drive allele and the NHEJ-derived resistance alleles (r2) reached equilibrium at approximately 43% each within 30 generations. We expected purification to occur exclusively in male gametes, based on *KLHL10*’s only previously identified function in male gametogenesis^13^ (Fig 4B). To our surprise, about 83% of frameshift mutations were eliminated in female offspring, since only one of the six frameshift mutations identified in female ovaries was passed onto offspring (Fig 4C). Lastly, we looked at the genotype composition dynamics under high purification. We observe progressive waves of the top three genotypes, with D/WT heterozygotes peaking around generation 15 as drive-carriers mate with wild-type individuals, then decline as the population transitions to predominately D/D homozygotes by generation 30. Targeting *KLHL10* outperforms targeting a haploinsufficient lethal gene as well as a recessive lethal gene (Supplemental Fig 4).

**Fig. 4:**
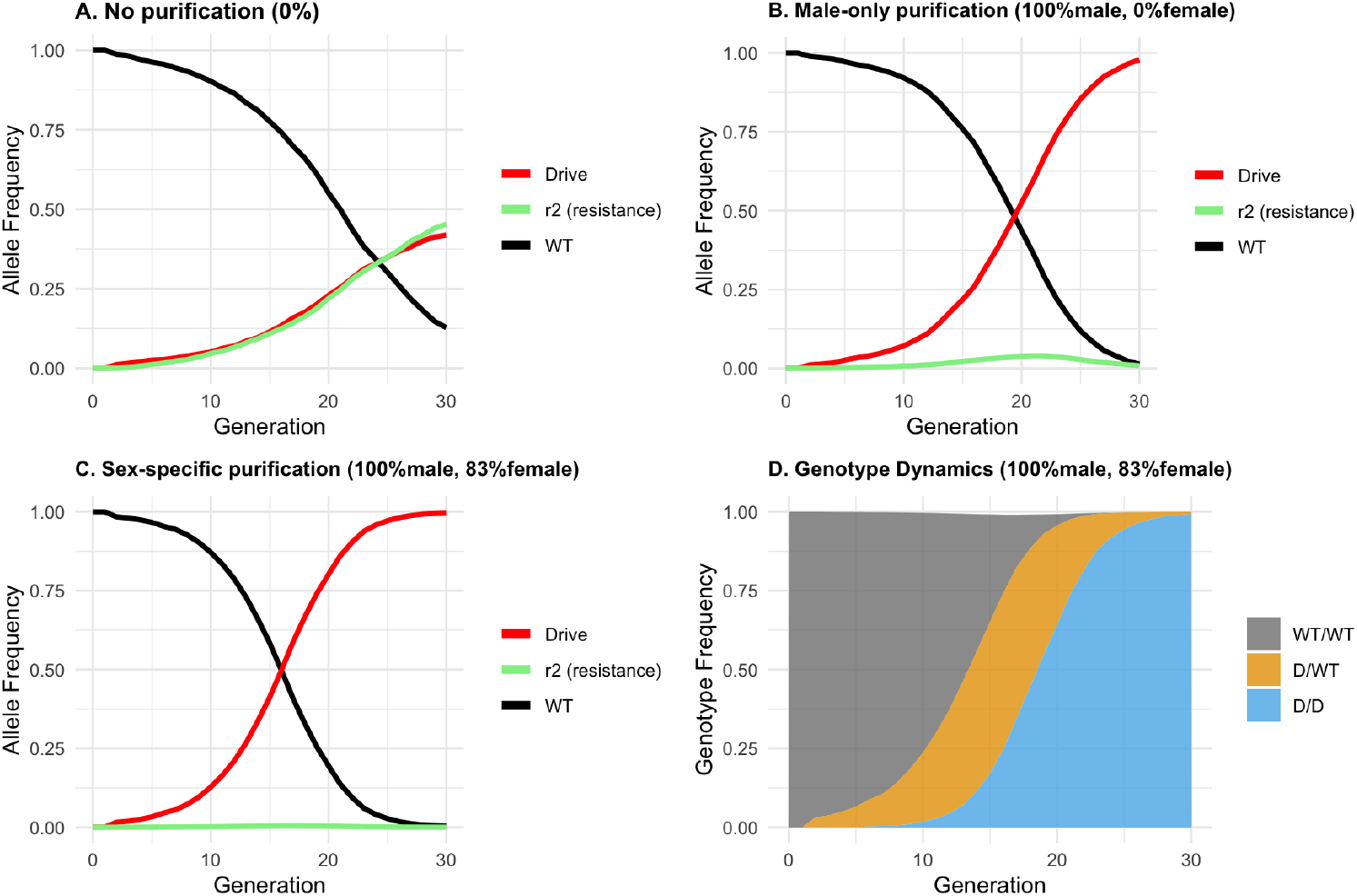
Modelling this system indicates enhanced purification of resistant alleles and drive efficiency. A.) No purification of NHEJ-derived resistance alleles, B.) Male-only purification (100%) and C.) Male (100%) and female (83%) purification. D.) Genotype dynamics over time under purification in males and females for the predominant three genotypes.

These modeling results demonstrate that purification of NHEJ-derived resistance alleles is critical for successful gene drive propagation given modest homing rates in rodents.

The model predicts that our experimental approach of targeting haploinsufficient *KLHL10* should provide sufficient purifying selection to enable drive spread, provided purification efficiency can be maximized through optimal guide selection and targeting.

## Discussion

This work provides a proof of principle for targeting a haploinsufficient spermatogenesis gene as a means of purifying undesired repair outcomes, suggests previously unknown functions of *KLHL10* in female fertility or oocyte development, and characterizes the effectiveness of LbCas12a in a mammalian gene drive context.

It is important to delineate what this study demonstrates from what remains to be shown. We validated that frameshift mutations at *KLHL10* are effectively excluded from male reproduction: a male with high frameshift mosaicism (63% somatic/91% germline) was infertile, while the coding sequence cassette fully rescued fertility in heterozygous and homozygous knock-in males. This confirms that the purification mechanism central to our proposed drive strategy, the elimination of NHEJ outcomes via spermatogenic selection, functions as predicted. However, because constitutive Cas12a expression precludes HDR, we did not observe or measure drive-mediated gene conversion. A full demonstration of this gene drive system will require germline-restricted expression during meiosis to enable HDR while maintaining the spermatogenic selection we characterize here.

We were surprised to find a potential implication of *KLHL10* in female fertility/embryogenesis when previously *KLHL10* has only been associated with fertility in males^13^. *KLHL10* belongs to the kelch-repeat-containing family, and proteins in this family are known to have varied roles including cellular communication, ubiquitination, and activities involved in cellular structure^20,21^. Of note, in *Drosophila*, kelch has a role in maintaining the actin organization of developing oocytes to facilitate cytoplasmic entry^21^. Additionally, mouse embryonic transcriptomic analyses have identified that *KLHL10* is expressed at the 8-cell and blastocyst stages^22^. Further research could examine the potential for this protein to have a role in mouse development, and could further elucidate the candidacy of this gene for gene drive technology.

Notably, male and female selection against *KLHL10* frameshifts appear to operate through distinct mechanisms with differing stringency. Male selection is absolute: high frameshift loads result in infertility, consistent with the established haploinsufficiency of *KLHL10* in spermatogenesis. Female selection, while strong, permits some frameshift inheritance—including the -8 deletion, which was tolerated in females but would presumably contribute to male infertility. This asymmetry suggests that *KLHL10* may play a less dosage-sensitive role in oogenesis or early embryogenesis than in spermatogenesis, or that selection occurs at a developmental stage where mosaicism allows some affected cells to be compensated by unaffected neighbors. Regardless of mechanism, the critical observation for gene drive applications is that male purification—the primary bottleneck this strategy addresses—appears to be complete.

We assume functional sperm remain sufficiently abundant after purification to maintain fertility^23,24^. This assumption warrants scrutiny: if sperm function requires a minimum threshold of viable cells, high cutting efficiency combined with low male HDR rates could compromise fertility. Our high-mosaicism infertile male (∼91% germline knockout) is consistent with this concern, though it is equally consistent with simple haploinsufficiency of individual sperm. However, mice are polygynous and subject to sperm competition, which should favor substantial overproduction of sperm relative to the minimum required for fertilization^25^. The extent to which this expected redundancy buffers against the loss of NHEJ-affected sperm remains unclear and may influence optimal drive parameters in practice.

We selected LbCas12a for its theoretical advantages: alternative PAM requirements, reduced recombination risk from scaffold-based array processing, and potential for iterative editing at previously cut sites^26,27^. However, our results do not support its use in mammalian gene drives. The experimental mutant scaffold failed to produce detectable cutting at its target site. Overall editing efficiency was moderate (38% of offspring mosaic), consistent with recent Cas12a mouse studies but substantially lower than published SpCas9 systems in comparable contexts^9–11,28^. One offspring showed possible evidence of iterative editing atop a parental +3 indel (Supplemental Figure 5), but this theoretical advantage does not compensate for reduced cutting efficiency. We recommend future mammalian gene drive designs prioritize SpCas9 or similarly robust nucleases.

Our results indicate that targeting haploinsufficient reproductive genes can provide strong selection pressure against NHEJ outcomes in both sexes. The selection pressure against NHEJ outcomes arising from targeting a haploinsufficient spermatogenesis gene is comparable to one targeting a haploinsufficient essential gene and stronger than targeting a recessive lethal gene (Supplemental Figure 4). It should be noted that the target site location may have influenced our results. PAM requirements for Cas12a led us to target exon 3, an area we have identified as a loop in a series of beta sheets. Due to this target site, it is possible that our system was more permissible to in-frame mutations. Targeting a structural section of the *KLHL10* beta barrel sheets, especially upstream sequences, could provide stronger selection against in-frame mutations. The mechanism by which the -8 frameshift escapes female selection remains unclear; multiplexed guides targeting distinct sites could provide redundancy against such allele-specific exceptions.

## Conclusion

In sum, we validate the core mechanism of a gene drive strategy targeting haploinsufficient spermatogenesis genes: frameshift mutations at *KLHL10* are excluded from male reproduction, while the coding sequence cassette rescues gene function without detectable fitness cost. Unexpectedly, female reproduction also selected against most frameshift alleles, suggesting additional developmental functions for *KLHL10*.

These results support further development of this approach using germline-restricted CRISPR expression to enable efficient HDR-mediated gene drive.

## Methods

### Mouse Model Generation and Husbandry

Transgenic mice were generated via CRISPR/Cas-mediated pronuclear injection by the Division of Comparative Medicine MIT Core using donor plasmids. For the LbCas12a mice, SpCas9 RNP was generated from commercial SpCas9 protein obtained from Integrated DNA Technologies (Coralville, IA, USA) and Rosa26 targeting crRNA (5’-ACTCCAGTCTTTCTAGAAGA-3’) from Synthego (Redwood City, CA, USA). The donor plasmid contained Rosa26 homology arms of 820bp, upstream, and 530bp, downstream, flanking a knock-in cassette of pCAG-NLS-LbCas12a-RR-NLS-βA(polyA), pGK-NLS-EGFP-NLS-βGH(polyA). For the Klhl10^sgRNA^ mice, LbCas12a RNP was generated from purified LbCas12a-RR obtained from the QB3 MacroLab at UC Berkeley (Berkeley, CA, USA) and the Klhl10-targeting crRNA (5’-GCCACCAATAGCGAATAGGATGG-3’) from Synthego (Redwood City, CA, USA). The donor plasmid contained Klhl10 homology arms of 500bp, upstream, and 912bp, downstream, flanking a knock-in cassette of Klhl10(CDS)-wtDR-crRNA1-mtDR1-crRNA2-mtDR2. The sequences for these elements is as follows: wtDR: 5’-TAATTTCTACTAAGTGTAGAT-3’, crRNA1: 5’-GCAGTGTTGAGACGCACGTAGCC-3’, mtDR1: 5’-TAATTTCTACTTAGTGTAGAT-3’, crRNA2: 5’-GCCACCAATAGCGAATAGGATGG-3’, mtDR2: 5’-TAATTTCTACTAATAGTAGAT-3’. Injected embryos were implanted into pseudopregnant females. Of 39 pups born from the Rosa26-LbCas12a injection, one pup carried the full length knock-in. Of the 22 pups born from the Klhl10-Klhl10^sgRNA^ injection, three pups carried the full length knock-in. Additional sequencing confirmed correct integration at the target locus in one mouse. Mice were generated and maintained on a C57BL/6NTac background. Food and water were available *ad libitum*. One mouse was determined to be infertile after harem breeding for 17 weeks produced no offspring.

### DNA Extraction and Genotyping

Genomic DNA was extracted from mouse ear tissue by overnight incubation of 12.5µL of a 20mg/mL stock proteinase K and 500uL of ‘tail buffer’ (10mM 1M Tris pH 8.0, 100mM 5M NaCl, 0.5M EDTA pH 8.0, 0.5% SDS) at 55C in 1.7mL microcentrifuge tubes. After overnight incubation, 167µL of 5M NaCl was added and mixed for 5 minutes, then centrifuged on a benchtop centrifuge at 16,000 x *g* for 10 minutes. 500µL of supernatant was removed and replaced with 500µL of isopropanol and inverted until a precipitate formed. DNA was then extracted using phenol-chloroform, and purified via ethanol precipitation.

For F3, mice were initially genotyped using Transnetyx (Cordova, TN) to determine which were homozygous or heterozygous for the engineered allele. For mice that were determined to be heterozygous, a followup DNA extraction, PCR, and sequencing was used to determine genotype.

### *KLHL10* Sanger and NGS Sequencing

For Sanger sequencing and Synthego ICE analysis, a pair of primers (oWK1307: 5’-TACCACCTCCACAAAACAAAACCATCTAT-3’, oWK1308:

5’-GAACTGTCTCAAATAAGTAAATGGGGGCT-3’) were used to amplify the targeted gDNA fragment with Primestar GXL DNA polymerase (Takara Bio USA San Jose, CA). PCR products were purified and Sanger sequenced using primer pair (oWK1414:

5’-TATCAAAAATACAGTATAGCAATTAACAGTG-3’, oWK1149:

5’-AAGCAAACTAAGGGGCTGG-3’). For NGS analysis, an adaptor-ligated amplicon was generated using primer pair (oWK1837-R1:

5’-TCCCTACACGACGCTCTTCCGATCTGTTAAGCGTTTTGACCCTGTGAAGAAAAC-3’, oWK1838-R2:

5’-GTTCAGACGTGTGCTCTTCCGATCTCCTGGTTCTTTACCTTCCCATAGAGTGTT-3’) and was sent to Quintara Bio (Cambridge, MA) for Illumina MiSeq sequencing. Fastq reads were then analysed using CRISPResso2 using default settings for paired end reads. For both Synthego ICE and CRISPResso2 analysis, we analyzed indels from the LbCas12-RR guide sequence (5’-GCAGTGTTGAGACGCACGTAGCC-3’) While the wildtype scaffold showed the expected activity, the experimental scaffold failed to produce detectable cutting at its target site *in vivo*. Downstream LbCas12a cutting analyses were therefore only conducted using the wildtype scaffolded guide.

### Population Modelling

Gene drive dynamics were simulated using a discrete-generation Wright-Fisher model with random mating, a carrying capacity of 10,000, and a survival rate of 0.95 per generation. Drive parameters and purification efficiencies are described in Results. Simulations were run for 30 generations with 100 D/D individuals introduced at generation 1. All modeling was performed in R (v4.4.1).

### Institutional ethics and regulatory approval

This work has obtained regulatory approval from the relevant institutions. All activities involving recombinant DNA and biosafety were carried out in accordance with guidelines set forth by the MIT Environment, Health, Safety (EHS) committee. Furthermore, all animal-related work adhered to the guidelines established by the MIT Institutional Animal Care and Use Committee (IACUC), following approved protocols (Protocol 2304000512)

