## Supplemental Material for "Increasing the effective gene drive homing rate by targeting the haploinsufficient spermatogenesis gene *KLHL10*"

| **Breeding pair (Parent of interest ID)** | **# of pups homozygotic for KLHL10 engineered allele** | **# of pups heterozygotic for KLHL10 engineered allele** | **# of het pups genotyped for indels** | **Parent somatic mosaic genotype (Sanger)** | **Offspring genotype** | **Ovary/ Testis germline mosaic genotype (NGS)** |
|---|---|---|---|---|---|---|
| 97<br>(3M) | 14 | 14 | 14 | 52% indel<br>(-8, -11) | WT: 14 | |
| 130<br>(659M) | 9 | 10 | 10 | 4% indel<br>(+14) | WT: 9<br>mosaic: 1 | +13: 3%<br>-6: 2%<br>WT: 94% |
| 132<br>(704M) | 0 | 0 | 0 | 63% indel<br>(+1, -11, -17) | NA | +1: 47%<br>-11: 22%<br>-17: 21%<br>WT: 9% |
| 98<br>(94F) | 9 | 15 | 14 | 21% indel<br>(-3,-7) | -3: 7<br>WT: 7 | |
| 123<br>(656F) | 14 | 16 | 16 | 9% indel<br>(-4) | +3: 6<br>WT: 10 | -4: 4%<br>+3: 2%<br>WT: 92% |
| 128<br>(674F) | 13 | 15 | 14 | 45% indel<br>(-8, +3) | +3: 7<br>-8: 3<br>WT: 3<br>mosaic: 1 | -8: 28%<br>+3: 11%<br>WT: 59% |
| 127<br>(705F) | 15 | 11 | 11 | 87% indel<br>(-6) | -6: 10<br>WT: 1 | -6: 82%<br>-11: 2%<br>+1: 2%<br>-17: 2%<br>WT: 12% |

Supplemental Table 1: Descriptions of the families analyzed in this experiment. 'WT' in 'Offspring Genotype' refers to offspring who inherited a *KLHL10* allele without indels, but these offspring were heterozygous for the engineered construct. Some mice were indicated to have more than one of an individual indel via trace decomposition analysis (ie: two different -4 deletions), for simplicity these are collapsed here but are considered different mutations during 'frequency' analysis. NGS %s ≤1% were removed from the table. Mouse 704 in breeding pair

132 was determined to be infertile after being kept with a harem of 3 female mice for 17 weeks without producing any offspring.

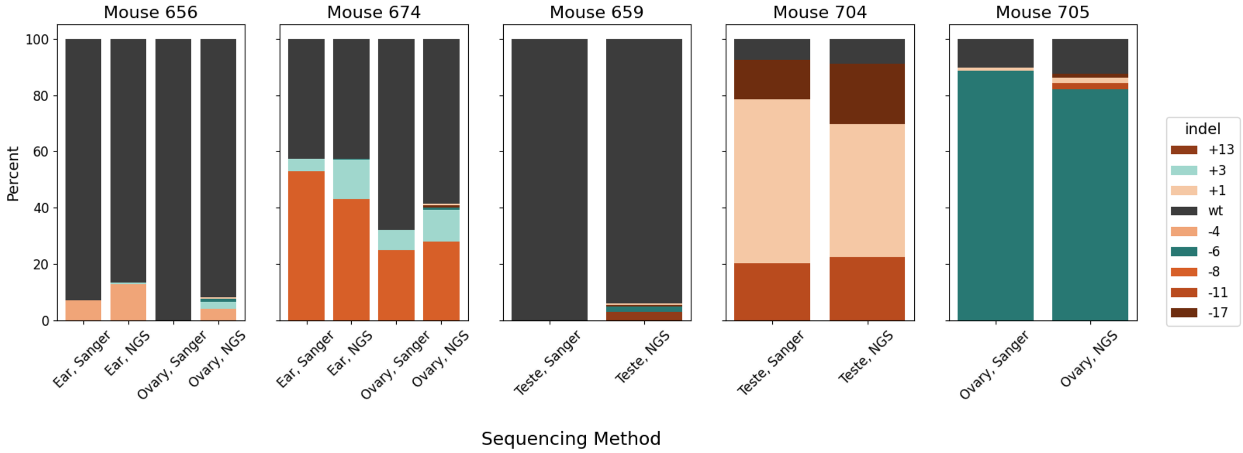


Supplemental Figure 1: Comparing sanger sequencing/ICE analysis to the NGS/CRISPResso analysis of mosaicism on the same tissue samples.

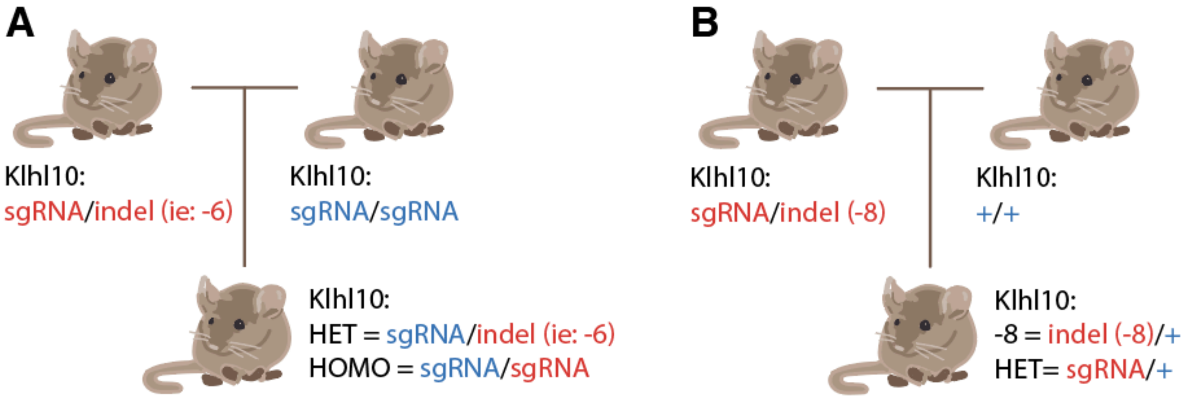


| Father (-6) | BP 145 | (n=12) | |
|---|---|---|---|
| Male | | Female | |
| HET | HOMO | HET | HOMO |
| 3 | 2 | 5 | 2 |

| Mother (-6) | BP 146, 147 | (n=17) | |
|---|---|---|---|
| Male | | Female | |
| HET | HOMO | HET | HOMO |
| 3 | 5 | 4 | 5 |

| Father (+3) | BP 141, 142 | (n=15) | |
|---|---|---|---|
| Male | | Female | |
| HET | HOMO | HET | HOMO |
| 4 | 3 | 5 | 3 |

| Mother (+3) | BP 143, 144 | (n=10) | |
|---|---|---|---|
| Male | | Female | |
| HET | HOMO | HET | HOMO |
| 3 | 1 | 1 | 5 |

| Mother (-8) | BP 140 | (n=6) | |
|---|---|---|---|
| Male | | Female | |
| -8 | HET | -8 | HET |
| 2 | 2 | 1 | 1 |

Supplemental Figure 2: Crossing frameshift and in-frame heterozygotes. BP = Breeding Pair, numbers indicate number of offspring in each category. A.) -6 and +3 *KLHL10* heterozygotes were fertile and no inheritance bias of the engineered allele was detected. B.) -8 *KLHL10* heterozygote female is fertile and did not produce any inheritance bias.

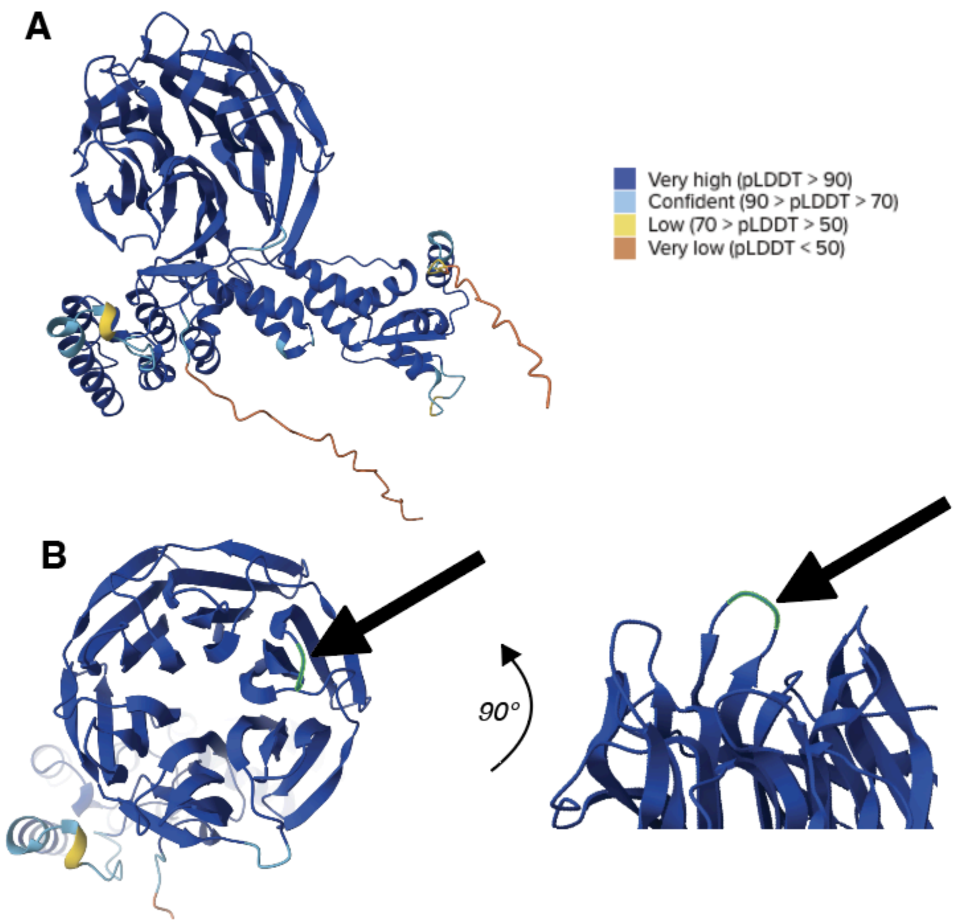


Supplemental Figure 3: Predicted cutting site. A.) Alphafold prediction of *KLHL10* protein structure. B.) Site of protein targeted for endonuclease cutting highlighted in green and indicated with an arrow. Green selection is 3 amino acids long.

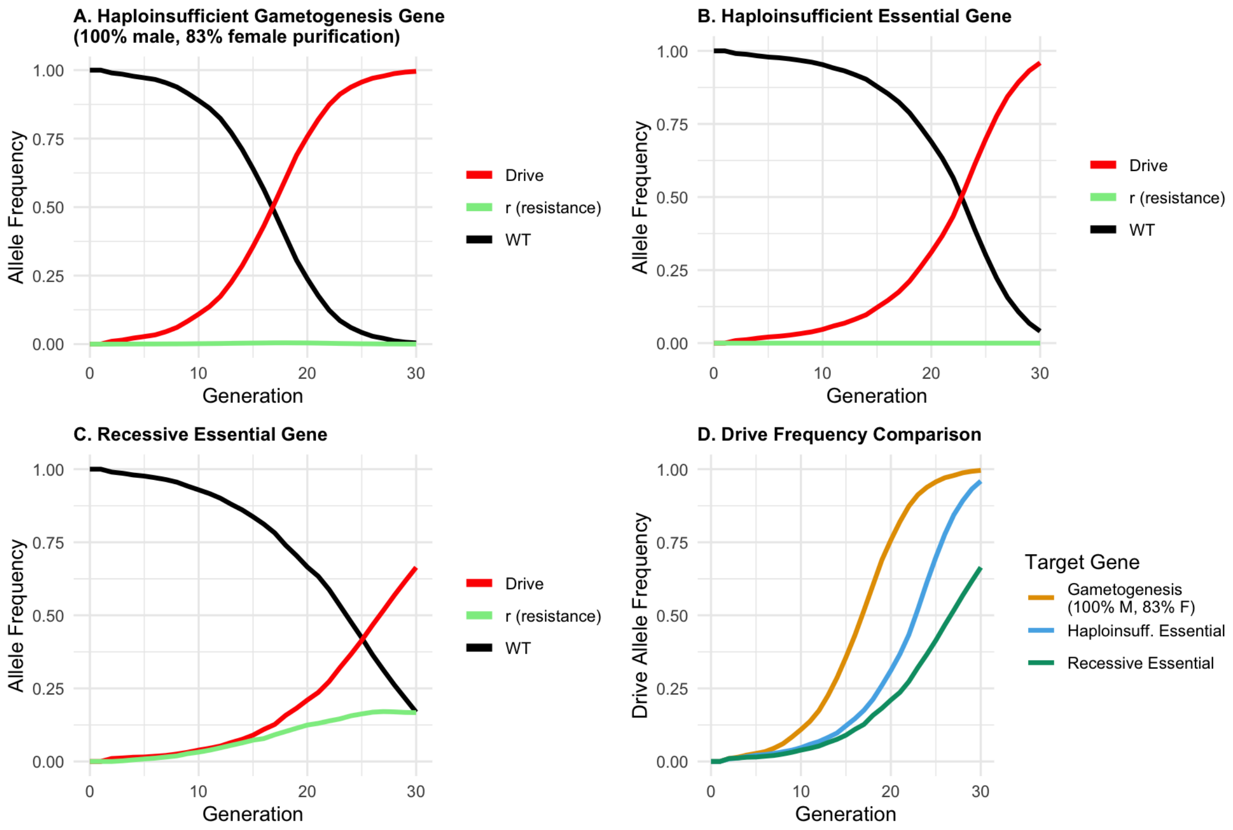


Supplemental Figure 4: Comparison of gene drive targeting strategies**.** Allele frequency dynamics for drives targeting (A) haploinsufficient gametogenesis gene Klhl10, (B) haplolethal essential genes (post-implantation lethality), and (C) recessive essential genes. (D) Drive frequency comparison. One-time release of 100 D/D individuals into a 10,000 wild-type population; sex-specific homing rates: 37.6% (female), 3.0% (male).

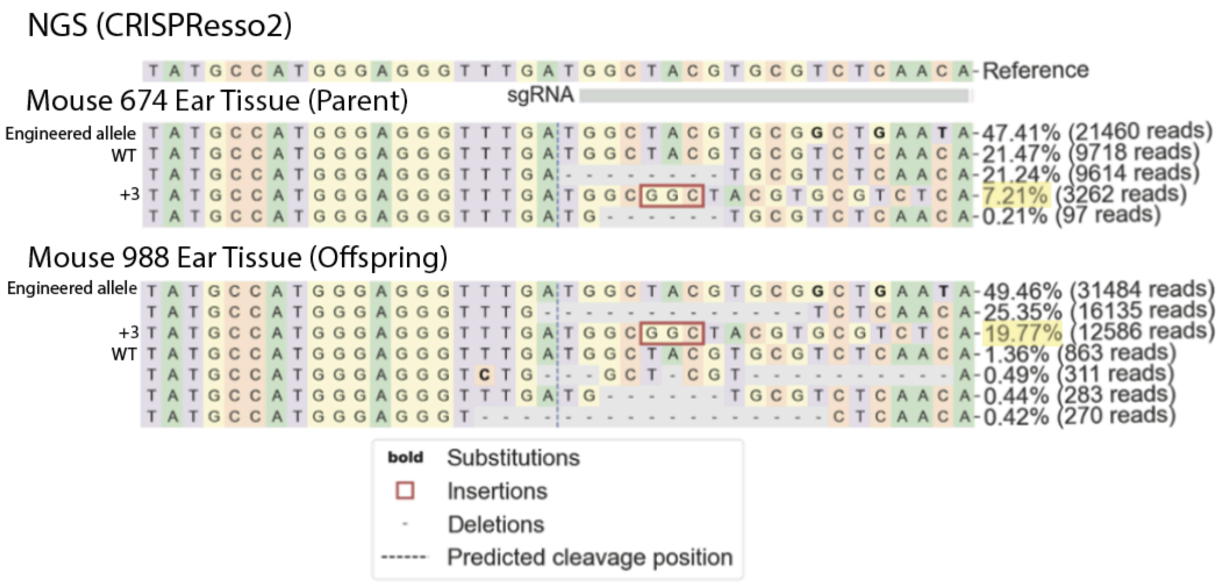


Supplemental Figure 5: Potential evidence for iterative Cas12 editing. Here, we have a parent mouse 674 that contains several mutations, one of which is a +3 insertion. One of the offspring of these mice had the same +3 insertion at a high rate(highlighted in yellow). Additionally, mouse 988 had few WT reads, suggesting that this mouse may have inherited the +3 indel and then underwent LbCas12 editing on top of that indel.
